# Changing expression system alters oligomerization and proinflammatory activity of recombinant human S100A9

**DOI:** 10.1101/2024.08.14.608001

**Authors:** Lauren O. Chisholm, Chae Kyung Jeon, James S. Prell, Michael J. Harms

## Abstract

S100A9 is a Damage Associated Molecular Pattern (DAMP) that activates the innate immune system via Toll-like receptor 4 (TLR4). Despite many years of study, the mechanism of activation remains unknown. To date, much of the biochemical characterization of S100A9 has been performed using recombinant S100A9 expressed in *E. coli* (S100A9^ec^). TLR4 is the canonical receptor for LPS, a molecule found in the outer membrane of *E. coli*, raising the possibility of artifacts due to LPS contamination. Here we report characterization of LPS-free recombinant S100A9 expressed in insect cells (S100A9^in^). We show that S100A9^in^ does not activate TLR4. This difference does not appear to be due to LPS contamination, protein misfolding, purification artifacts, or differences in phosphorylation. We show instead that S100A9^in^ adopts an altered oligomeric state compared to S100A9^ec^. Disrupting oligomer formation with the *E. coli* disaggregase SlyD restores activity to S100A9^in^. Our results also indicate that the oligomeric state of S100A9 is a major factor in its ability to activate TLR4 and that this can be altered in unexpected ways by the recombinant expression system used to produce the protein.

## INTRODUCTION

S100A9 is a small, dimeric calcium binding protein that is highly abundant in neutrophils^1^. S100A9 is a Damage Associated Molecular Pattern (DAMP) that activates inflammation via Toll-like receptor 4 (TLR4)^2–5^. S100A9 also forms a heterodimer with S100A8 known as calprotectin, an anti-microbial protein functioning in nutritional immunity^6^. As a DAMP, S100A9 activates the immune receptor TLR4 by an unknown mechanism. This activity has been demonstrated many times, in many different systems^2–5,7–10^, and has been associated with negative outcomes in neurodegenerative diseases^11,12^ and many cancers^13–15^.

One challenge for studies of S100A9 activity is that its receptor, TLR4, is the canonical receptor for lipopolysaccharide (LPS), a component of the outer membrane of gram-negative bacteria^16–18^. LPS is a common contaminant in recombinant proteins purified from *E. coli*, potentially leading to spurious activation of TLR4. Most studies of DAMPs activating TLR4 have used proteins expressed in gram-negative bacteria^2,4,5,19^, followed by purification steps to remove LPS. Because of this, some have suggested that DAMP activation of TLR4 is nothing more than an experimental artifact^20^.

We decided to remove LPS contamination at the source and recombinantly express S100A9 in eukaryotic cells. We show that, while S100A9 prepped out of *E. coli* (S100A9^ec^) activates TLR4, S100A9 prepped out of insect cells (S100A9^in^) does not. We rule out protein misfolding, post-translational modification, and simple LPS contamination as causes for the difference in activity. We show that S100A9^in^ displays altered oligomeric states compared to S100A9^ec^. We find disrupting oligomer formation of S100A9^in^ using an *E. coli* disaggregase restores proinflammatory activity. This suggests that S100A9 can indeed activate TLR4; however, its oligomeric state is a key determinant of proinflammatory activity.

## RESULTS AND DISCUSSION

### Changing S100A9 expression system changes proinflammatory activity

We set out to purify human S100A9 from several cell types to determine the extent to which S100A9 un-contaminated with LPS could activate TLR4. We recombinantly expressed human S100A9 from three cell types: Rosetta BL21(DE3) pLysS *E. coli* (S100A9^ec^), HighFive insect cells (S100A9^in^), and HEK293F human cells (S100A9^hek^). We were unable to purify S100A9^hek^ due to extensive proteolysis of S100A9 by human cells^21,22^. S100A9^ec^ and S100A9^in^ were readily expressed and purified to >99% purity by SDS-PAGE, so we focused on a comparison between these two recombinant proteins. For S100A9^ec^, we purified the protein with 3 chromatography steps (Ni-NTA, followed by sequential anion exchange at pH 8, then pH 6), followed by removal of LPS with an endotoxin removal kit. For S100A9^in^, we achieved high purity in a single Ni-NTA step.

To measure TLR4 activity, we used a previously established activity assay^10,23,24^. In this assay, we transiently transfect HEK293T cells with plasmids encoding TLR4 and its co-receptors MD-2 and CD14, then measure TLR4 activity with a firefly luciferase behind an NF-κB promoter. In this way we can test the TLR4-specific proinflammatory activity of various agonists. To control for possible LPS contamination of recombinantly prepared proteins, we can include polymyxin B (PB), which binds to LPS and prevents LPS activation of TLR4 (Fig 1A)^2,10,24^.

**Figure 1:**
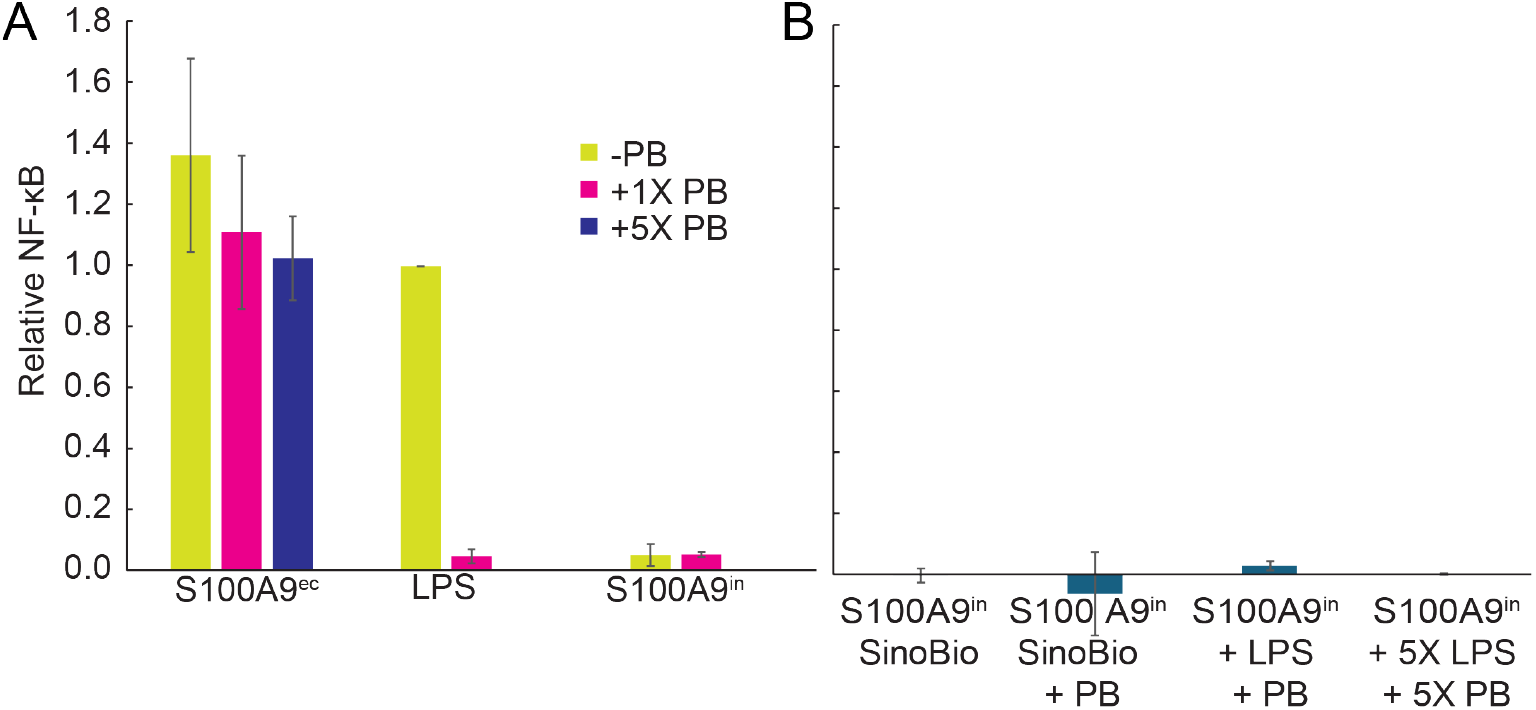
S100A9^in^ does not activate TLR4, and does not deliver LPS to TLR4. A) Relative NF-κB in response to S100A9^ec^ with multiple PB concentrations (0X, 1X and 5X), LPS with and without PB, and S100A9^in^ with and without PB. NF-κB output is normalized such that the reponse to LPS = 1.0. Bar color indicates amount of PB added. B) Relative NF-κB in response to S100A9^in^ that has been pre-incubated with LPS, (followed by the addition of PB), and to S100A9^in^ purchased from Sino Biological. For 1X: LPS concentration is 200ng/mL, 1X PB concentration is 200ug/mL, S100A9 concentration is 2uM. Data displayed is the average of 3+ biological replicates, error bars indicate standard error of the mean (SEM).

We first tested S100A9^ec^, finding it activates TLR4 even with large amounts of polymyxin B (PB) (Fig 1A). This matches previous reports^2,5,10,25^. We next tested S100A9^in^. To our surprise, it did not activate TLR4 (Fig 1A). To rule out a problem with our purification technique, we purchased and tested commercially available S100A9^in^ (Sino Biological). Like our prepared protein, commercial S100A9^in^ did not activate TLR4 (Fig 1B).

### S100A9^ec^ does not deliver LPS

We hypothesized that S100A9^ec^ was activating TLR4 by delivering LPS to TLR4, rather than S100A9 directly activating TLR4. To test this, we mimicked LPS contamination by pre-incubating S100A9^in^ with purified LPS. We then treated this sample with PB—in the same way we treated S100A9^ec^—and tested its activity. This treatment condition was also inactive (Fig 1B), ruling out LPS delivery by S100A9.

### The difference is not due to a post-translational modification

We next hypothesized the difference was due to a post-translational modification of S100A9^in^ not present in S100A9^ec^. S100A9 has one well characterized post-translational modification – a phosphorylation at position T113^26–28^. This has been shown to modulate certain activities of S100A9. For example phosphorylation inhibits S100A9-induced polymerization of tubulin^28^. It has also been reported that phosphorylation modulates the inflammatory state of calprotectin^29^. To assess whether S100A9^in^ might be phosphorylated at position T113, we performed western blots against the protein using either a generic anti-S100A9 antibody (1C22 Abnova) or antibody specific to S100A9 T113-p (#12782 Signalway Antibody). We found that the generic antibody recognized both S100A9^ec^ and S100A9^in^ (Fig 2A), but that the phosphorylation-specific antibody only recognized S100A9^in^ (Fig 2A). We further validated the phosphorylation of T113 using top-down mass spectrometry (Oregon State University, Mass Spectrometry Core).

**Figure 2:**
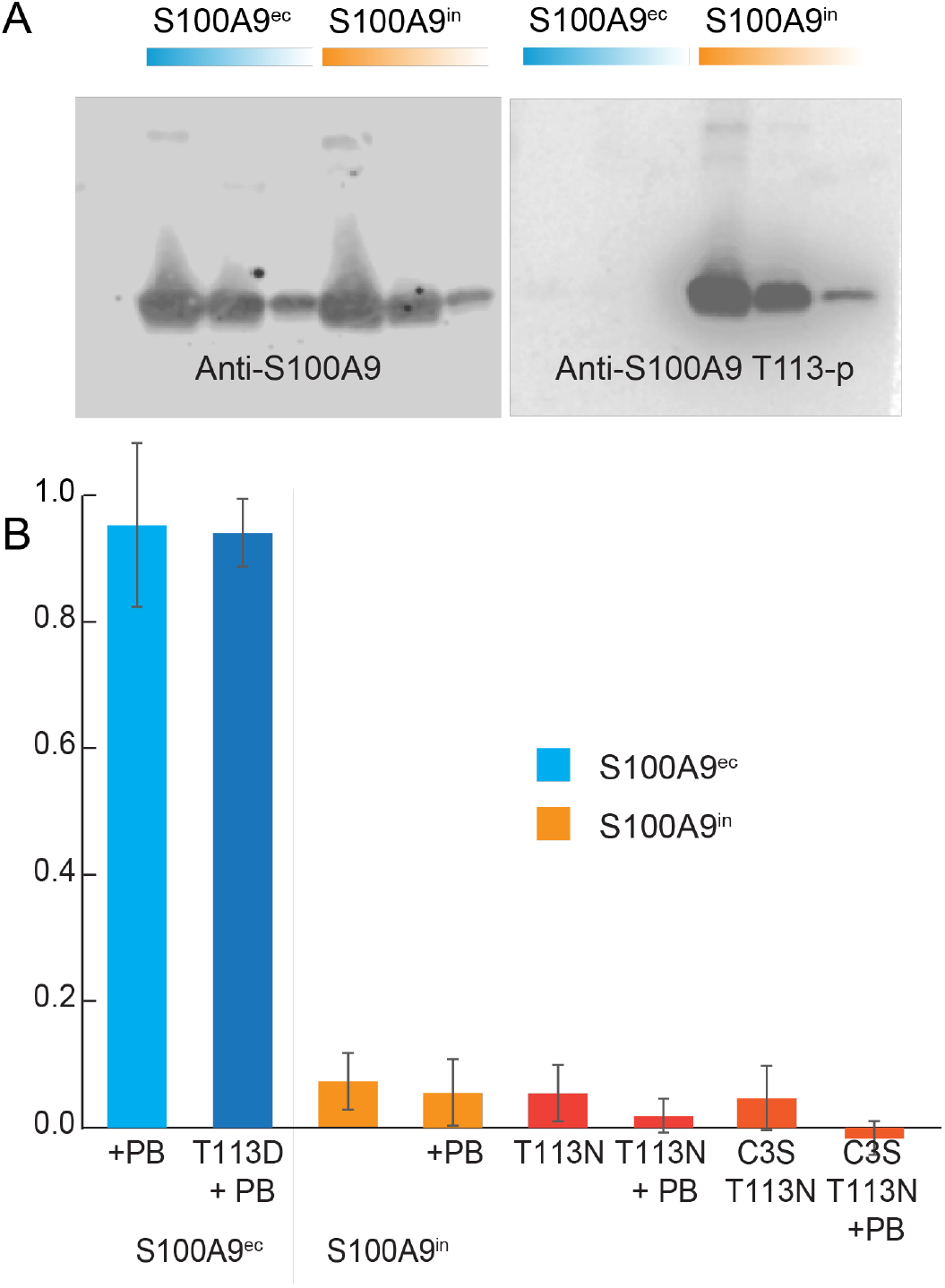
S100A9^in^ is phosphorylated, but this does not alter activity. A) western blot of S100A9^ec^ and S100A9^in^ with anti-S100A9 antibody (1C22 Abnova) and anti-S100A9 T113-p antibody (#12782 Signalway Antibody). The western blot images were cropped to exclude unnecessary white space, and converted to greyscale for improved visibility. B) Relative NF-κB in response to T113 mutations to S100A9^in^ and S100A9^ec^. NF-κB is normalized such that wildtype S100A9^ec^ response = 1.0. PB concentration is 200ug/mL, S100A9 concentration is 2uM. Data displayed is the average of 3+ biological replicates, error bars indicate standard error of the mean (SEM).

We hypothesized that phosphorylation was somehow modulating TLR4 activation. To test this hypothesis, we first attempted to dephosphorylate S100A9^in^ by digesting with commercially available calf intestinal alkaline phosphatase (CIP – New England Biolabs). Our efforts at dephosphorylation were, however, unsuccessful as assessed by both western blot and MALDI-TOF mass spectrometry. We then turned to site-directed mutagenesis of S100A9^in^ and S100A9^ec^. To remove phosphorylation in the insect cell protein, we introduced S100A9^in^ T113N; to mimic phosphorylation in the *E. coli* protein, we introduced S100A9^ec^ T113D.

Neither removing the phosphorylation nor introducing a phosphomimetic had any effect: S100A9^in^ T113N did not restore activity to S100A9^in^ (Fig 2B), while S100A9^ec^ T113D did not disrupt TLR4 activity from wildtype S100A9^ec^ (Fig 2B). We also tested S100A9^in^ T113N in a cysteine free background (C3S) (Fig 2B), to confirm that disulfide formation was not responsible for the difference in activity. These results suggest that some other feature of the protein, not the post-translational modification, differs between S100A9^in^ and S100A9^ec^.

### S100A9^in^ and S100A9^ec^ likely have different high-order structures

We set out to determine other differences between S100A9^ec^ and S100A9^in^ that might explain the difference in activity. We measured three spectra: far-UV circular dichroism (CD), near-UV CD, and intrinsic fluorescence. We made these measurements both in the presence and absence of calcium, as S100A9 is known to undergo a conformational change in response to calcium binding^30^.

Based on their far-UV CD spectra, both S100A9^ec^ and S100A9^in^ were primarily α-helical and responded similarly to the addition of calcium (Figure 3A). Likewise, we observed nearly identical intrinsic fluorescence spectra for both proteins (Figure 3B). The two purifications differed, however, in their near-UV CD spectra. S100A9^in^ exhibited a strong peak around 265 nm that was absent in S100A9^ec^. This suggested a difference in the tertiary or quaternary structures of the two proteins

**Figure 3.**
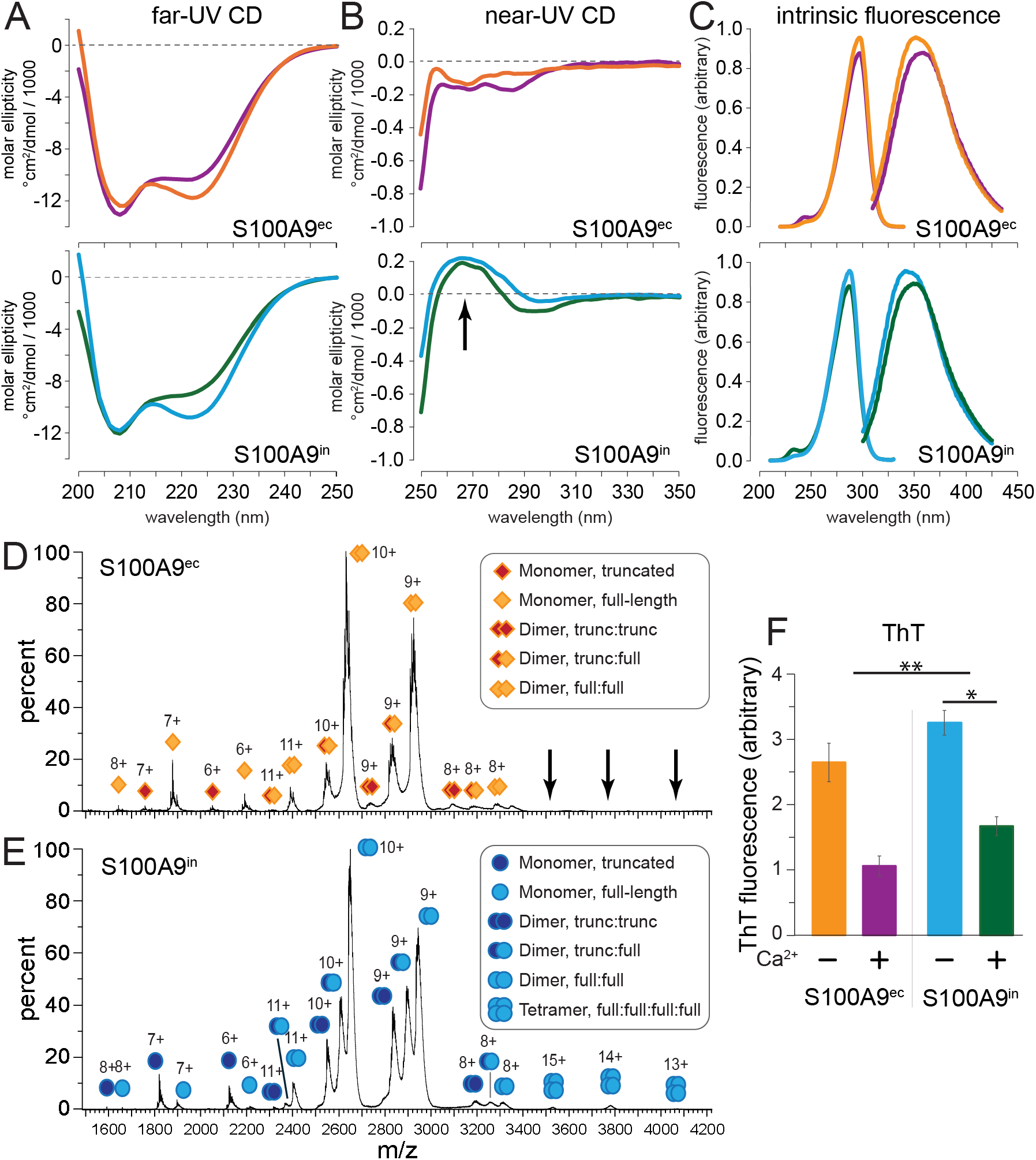
S100A9^in^ has a different quaternary structure than S100A9^ec^. Throughout the figure, orange indicates S100A9^ec^, purple indicates S100A9^ec^ + calcium, blue indicates S100A9^in^, and green indicates S100A9^in^ + calcium. A) Far-UV CD spectra of S100A9^ec^ (top) and S100A9^in^ (bottom). B) Near-UV CD spectra of S100A9^ec^ (top) and S100A9^in^ (bottom). The arrow indicates the peak present in S100A9^in^ but not S100A9^ec^. C) Intrinsic fluorescence emission and excitation spectra of S100A9^ec^ (top) and S100A9^in^ (bottom). Excitation spectra (emission at 345 nm) are on the left; emission spectra (excitation at 288 nm) are on the right. D) Native mass spectrum of S100A9^ec^. Inferred species are annotated as indicated in the key. (We observed full-length and truncated forms of S100A9, as indicated). The locations of the tetramer peaks observed for S100A9^in^ but not S100A9^ec^ are indicated with arrows. E) Native mass spectrum of S100A9^in^, annotated similarly to panel D. F) Amyloid formation as measured by ThT fluorescence (ex/em: 450/480) after an 8-hour incubation at 37 °C. Bars show representative results for one biological replicate; error bars are standard error of three technical replicates. P-values were calculated using a paired 2-tailed Student’s t-test on four biological replicates. ** P < 0.01; * P < 0.05

### The proteins have different oligomeric and aggregation behaviors

We attempted to determine the tertiary structure of S100A9^in^ using x-ray crystallography but were unable to obtain usable crystals. To probe the quaternary structure, we used native mass spectrometry (nMS) to measure the oligomeric states of S100A9^in^ and S100A9^ec^ as previously reported^31^. S100A9^in^ and S100A9^ec^ were measured at approximately the same bulk concentration. As expected, both proteins had peaks corresponding to dimers. S100A9^in^, however, also populated a tetrameric form (Fig 3D & E). Of all species detected by native MS, the prominence of dimer peaks in both S100A9^ec^ and S100A9^in^ suggests that dimer form of S100A9 are the relevant and preferable species in solution.

S100A9 is also known to form amyloid like fibrils^32,33^. Therefore, we hypothesized that S100A9^in^ may also be more prone to amyloid fibril formation. Using the amyloid fibril specific fluorescent dye, ThioflavinT (ThT), we measured the capacity of S100A9^in^ and S100A9^ec^ to form these fibrils, in the presence and absence of calcium. S100A9^in^ exhibited a significantly higher ThT fluorescence compared to S100A9^ec^ regardless of buffer conditions (p = 0.008). This difference could be due to a difference in total amount of fibrils formed, or in the structure of the fibril. The addition of calcium significantly reduced ThT fluorescence for S100A9^in^ (p = 0.035), consistent with previous reports on S100A9^ec 32^.

### Disrupting oligomer restores some activity

We next sought to understand why the two purifications gave different oligomeric states. We hypothesized the cause might be the bacterial protein SlyD, which binds weakly to Ni-NTA columns and can co-purify with recombinant proteins^34,35^. SlyD acts as a chaperone and disaggregase and could thus plausibly alter the oligomeric state and activity of recombinant S100A9. Further, because it is a chaperone, even a tiny amount of contaminant could lead to the result—consistent with the observation that S100A9^ec^ was >99% pure by SDS-PAGE.

We revisited our purification fractions and discovered that a protein with a molecular weight consistent with SlyD (∼25 kDa) eluted from our Ni-NTA column just prior to S100A9. We first established that this fraction (“HisA”), on its own, was not sufficient to activate TLR4 above background (P = 0.097) (Figure 4A). To test whether it could, however, increase the ability of S100A9^in^ to activate TLR4, we added this fraction to S100A9^in^. This led to a high NF-κB signal, even in the presence of large excess of PB to sequester LPS contamination (P = 0.0041) (Figure 4A). Having observed this change in activity, we next wanted to see if it corresponded to the predicted change in oligomeric state of S100A9. We added “HisA” to the protein and measured its native mass spectrum. All monomer and dimer species observed were comparable to the previous nMS of S100A9^in^; however, addition of HisA eliminated the tetramer peaks in the native mass spectrum (Fig C & D).

**Figure 4:**
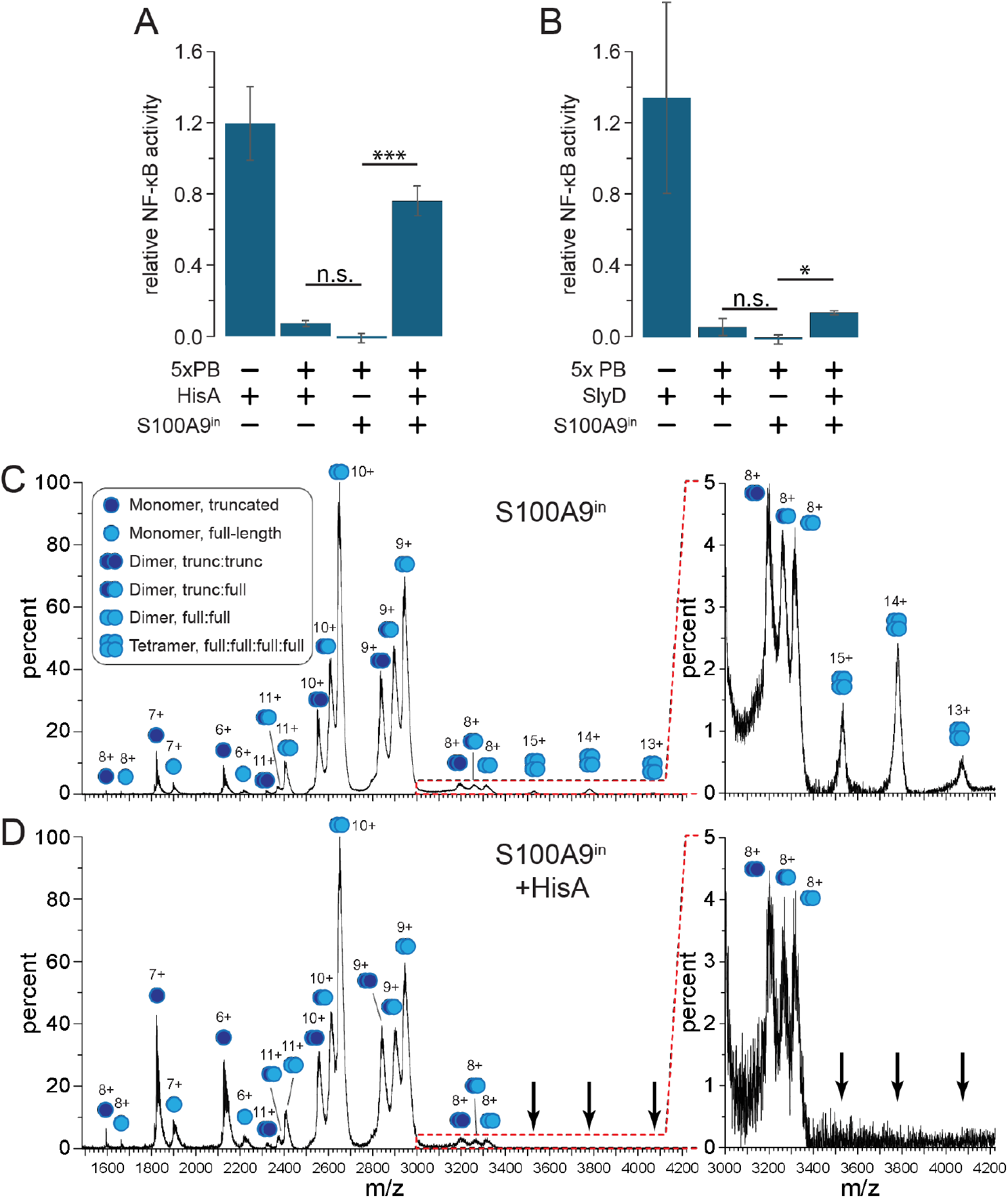
Bacterial protein contaminant SlyD alters S100A9 oligomeric state and activity. A) Purification fraction HisA restores some level of activity. B) Purified SlyD restores TLR4 activity, but not as potently as HisA. NF-κB is normalized such that Wildtype S100A9^ec^ response = 1.0. PB concentration is 200 μg/mL, S100A9 concentration is 2 μM. SlyD and HisA concentration is 0.2 μM. Data displayed is the average of 3+ biological replicates, error bars indicate standard error of the mean (SEM). P-values were calculated using a paired 2-tailed Student’s t-test. *** P < 0.005; * P < 0.05. C-D) Native mass spectra show disruption of tetramer with the addition of HisA fraction. Insets zoom in on tetramer region. Panel C shows the protein without the addition of HisA; panel D shows the protein after addition of HisA.

To test whether SlyD was cause of the change in activity, we tested the ability of commercially available SlyD (AbCam) to restore S100A9^in^’s ability to activate TLR4. Similar to HisA, SlyD alone does not activate TLR4 in the presence of high concentrations of PB (P = 0.38) (Figure 4B). When pre-incubated with S100A9^in^ followed by the addition of PB, S100A9^in^ exhibits activity above background (P = 0.015) (Figure 4B). This activity is less than that observed for HisA+S100A9^in^ (Figure 4A). This could be due to the concentration of SlyD in HisA differing from that in the purified sample, lower activity of the commercial preparation, or other factors in the HisA fraction contributing to the activity of the S100A9.

## CONCLUSIONS

We set out to purify LPS-free recombinant S100A9 using insect cells. We found that S100A9^in^ retained calcium binding and secondary structure but showed differences in tertiary structure and oligomeric state compared to S100A9^ec^. Most importantly, S100A9^in^ lacks S100A9’s reported proinflammatory activity. Our results also indicate that the oligomeric state of S100A9 is a major factor in its ability to activate the immune system. There are many open questions raised by our findings: What is the structure of S100A9^in^ and how does it differ from S100A9^ec^? How does either structure compare to endogenously expressed human S100A9?

Our findings highlight the difficulty intrinsic in mapping results with recombinant proteins to their biological context. Indeed, this is not the first time changing the expression system of a recombinant S100 has turned the field on its head – S100A3 was found to adopt different structure when expressed in insect cells rather than *E. coli* ^36^.

Our findings also emphasize the delicate balance the immune system must maintain: even small changes to the tertiary structure and oligomeric state of S100A9 completely abolished its proinflammatory activity. Learning more about the native structure and oligomeric state of S100A9 will be critical to better understanding—and maybe someday intelligently modulating— its activity in the innate immune system.

## METHODS

### Native Mass Spectrometry (nMS)

Protein samples (S100A9^ec^ and S100A9^in^) were buffer exchanged into 200 mM ammonium acetate at pH 7 using Micro Bio-spin P6 columns (Bio-Rad, Hercules, CA). Buffer exchanged samples were then diluted to a working concentration of 10 μM. HisA was added in 1:1 volume with S100A9^in^ immediately prior to the introduction to the instrument. These protein samples were analyzed with native MS on a Waters Synapt G2-S*i* quadrupole–ion mobility– time-of-flight instrument (Milford, MA). Samples were introduced using borosilicate capillary needles (prepared in-house with emitter i.d. ∼450 nm) threaded with a platinum wire to allow nano-electrospray ionization (nESI). The electrospray was operated at capillary voltage of 0.3-0.5 kV with the sample cone and temperature at 25 V and 25 °C, respectively, in positive mode. The Trap cell was operated with an argon gas flow of 10 mL/min. The Collision Energy for both the Trap and Transfer cells was set to 5 V. For ion mobility separations, the IMS cell was pressurized at ∼3.4 mbar nitrogen buffer gas. IM separation was performed with a traveling wave height of 30 V and wave velocity of 600 m/s. All native MS data were collected over the *m*/*z* range of 500-8000.

### *Recombinant S100A9 expression and purification from* E. coli

We expressed and purified S100A9 as previously described ^19,25^. Briefly, we expressed cysteine-free human S100A9 (C3S) in the pETDUET-1 vector in Rosetta BL21(DE3) pLysS *E. coli*. We purified S100A9 using three chromatography steps: immobilized metal ion affinity (HisTrap) at pH 7.4, anion exchange (HiTrap Q) at pH 8, followed by another anion exchange (HiTrap Q) at pH 6. We verified protein purity was >95% by SDS-PAGE. Proteins were stored at -80 °C until needed. We determined protein concentration using A_280_ with an extinction coefficient of 6990 M^-1^ cm^-1^ (monomer). The protein concentrations reported in this manuscript are μM dimer.

### Recombinant S100A9 expression in HighFive cells

Recombinant expression of S100A9 in insect cells was performed using an existing protocol^37^. Bacmid DNA was generated by cloning human S100A9 into the pFastBac1 plasmid, which was then transformed into DH10Bac cells. The human S100A9 gene was purchased from Genscript (Clone ID: OHu25452C Accession No.: NM_002965.4). pFastBac1 and DH10Bac cells were a gift from Scott Hansen. Bacmid DNA was then transfected into Sf9 cells. Baculovirus generated from this transfection was further expanded to obtain a suitable volume and viral titer for protein expression. The resulting baculovirus was used to infect 1L HighFive cells for protein purification. Sf9 and HighFive cells were a gift from Scott Hansen. HighFive cells were harvested using centrifugation, and lysed using a dounce. The lysate was treated with EDTA-free protease inhibitor cocktail (Sigma Aldrich). We purified the S100A9 using one chromatographic step. The lysate was incubated with 1 mL of Ni-NTA agarose (Thermo Fisher) at 4 °C for 1 hour. We then washed the resin twice with 25 mL of 25 mM Tris, 100 mM NaCl, 5mM BME, 0.1% Tween pH 7.4 containing 25 mM imidazole. We eluted protein with 10 mL 25 mM Tris, 100 mM NaCl, 5mM BME, 0.1% Tween pH 7.4 containing 500 mM imidazole. We verified protein purity was >95% by SDS-PAGE. We concentrated and buffer exchanged proteins into 25 mM Tris, 100 mM NaCl, 5mM BME, 0.1% Tween pH 7.4, then flash-froze dropwise into liquid nitrogen. Proteins were stored at -80 °C until needed. We determined protein concentration using A_280_ with an extinction coefficient of 6997 M^-1^ cm^-1^ (monomer). The protein concentrations reported in this manuscript are μM dimer.

### NF-κB activity assay

We measured NF-κB activity, and normalized the data as previously described^10,19,25,38^. In brief, we co-transfected plasmids individually encoding TLR4, MD-2, CD14, and an Nf-kB luciferase reporter into HEK293T cells seeded into a 96 well plate. After stimulation with either LPS or S100A9 we then lyse cells and measure luminescence using the Promega Dual-Glo Luciferase kit. For data processing and normalization between experiments, each plate contained the following four treatments: mock (PBS), 200 ng/mL LPS, 200 ng/mL LPS with 200 μg/mL polymyxin B, and 2 μM S100A9 with 200 μg/mL polymyxin B.

### Circular Dichroism

Circular dichroism and intrinsic fluorescence measurements were collected on a J-815 CD spectrometer. All samples were dialyzed O/N at 4* prior to measurement in 25 mM Tris, 100mM NaCl, 2mM TCEP, pH7.4. Near UV (250-350nm) CD and intrinsic fluorescence were collected at 25uM S100A9 (monomer), in a 1cm cuvette. Far UV (200-250nm) CD was collected at 10uM S100A9 (monomer) in a 1mm cuvette. Samples with calcium contained 1mM CaCl2, followed by the addition of 5mM EDTA for no calcium measurements. Buffer measurements were collected both with and without calcium. Near and Far UV CD spectra were collected 3 times. Fluorescence excitation spectra was collected at emission 345nm, emission was collected at excitation 288nm. Using Jasco’s software, the data was accumulated, buffer measurement was subtracted, and Savitsky-Golay filtered (level 9).

### Western Blotting

Blots were performed using either monoclonal mouse anti-S100A9 primary antibody, M13, clone 1C22 (Abnova) paired with IRDye 800CW Goat anti-Mouse IgG1 Secondary (Licor), or Polyclonal anti hS100A9 (Phospho-Thr113) Rabbit primary antibody #12782 (Signalway Antibody) paired with IRDye® 800CW Goat anti-Rabbit IgG Secondary Antibody (Licor).

### Top Down Mass Spectrometry

Top down mass spectrometry (TOF MS ES+ and FTMS) was performed by the mass spectrometry core at Oregon State University (data not shown).

### Thioflavin T Fluorescence

ThT fluorescence was measured as previously described^32^. Measurements were collected using 50uM ThT (Sigma Aldrich) and 50uM S100A9 (monomer), in a lidded Corning half area black plate after 8 hours of incubation, using an excitation/emission of 450/480. Buffer conditions: 50mM HEPES, pH7.4, 2mM TCEP, and EDTA-free protease inhibitor. Additionally, samples with calcium contained 1mM CaCl2, samples without calcium contained 1mM EDTA.

## ACKNOWLEDGEMENTS

We thank current and former members of the Harms lab for helpful discussion and input. We thank the Hansen lab for training in baculovirus expression and gift of the pFastBac1 plasmid, and Sf9 and HighFive cells. Funding: 5T32GM007759 (LOC), The John Keana Graduate Student Fellowship (LOC), NIGMS R01-GM146114 (MJH), and NIGMS R01-GM144507 (JSP).

